# Molecular evolution and developmental expression of melanin pathway genes in Lepidoptera

**DOI:** 10.1101/2020.03.11.987669

**Authors:** Muktai Kuwalekar, Riddhi Deshmukh, Ajay Padvi, Krushnamegh Kunte

## Abstract

Pigmentation is involved in a wide array of biological functions across insect orders, including body patterning, thermoregulation, and immunity. The melanin pathway, in particular, has been characterized in several species. However, molecular evolution of the genes involved in this pathway is poorly characterized, and their roles in pigmentation of early developmental stages are just beginning to be explored in non-model organisms. We traced the molecular evolution of six melanin pathway genes in 53 species of Lepidoptera covering butterflies and moths, and representing over 100 million years of diversification. We compared the rates of synonymous and nonsynonymous substitutions within and between these genes to study signatures of selection at the level of individual sites, genes, and branches of the gene tree. We found that molecular evolution of all six genes was governed by strong purifying selection. Yet, a number of sites showed signs of being under positive selection, including in the highly conserved domain regions of three genes. Further, we traced the expression of these genes across developmental stages, tissues, and sexes in the *Papilio polytes* butterfly using a developmental transcriptome dataset. We observed that the expression patterns of the genes in *P. polytes* largely reflected their known tissue-specific function in other species. The expression of sequentially acting genes in the melanin pathway was correlated. Interestingly, four out of six melanin pathway genes (*ebony, pale, aaNAT*, and *DDC*) showed a sexually dimorphic pattern of developmental heterochrony; i.e., females showed peak activity much earlier in pupal development compared to that of males. Our evolutionary and developmental analyses suggest that the vast diversity of wing patterning and pigmentation in Lepidoptera may have been aided largely by differential developmental regulation of genes in a highly conserved pathway, in which the sequence evolution of individual genes is highly constrained.

## INTRODUCTION

The evolution of color patterns contributes to the striking diversity of lifeforms. Coloration is a critical adaptation that impacts an organism’s intra- and inter-specific communication and interactions. In several cases, emergence of sexual dimorphism and polymorphism is also linked to the evolution and differential expression of pigmentation genes (Wittkopp et al., 2009; Miyazaki et al., 2014; Yassin et al., 2016). Among Lepidoptera, wing coloration and patterning are remarkably diverse, which have shaped several adaptations such as aposematism, crypsis, mimicry, and thermoregulation (True, 2003; Hegna et al., 2013; Olofsson et al., 2013; Kronforst and Papa, 2015; Nadeau, 2016; Deshmukh et al., 2018; van’t Hof et al., 2019). In insects, melanins, ommochromes, and pterins are three major biosynthesized pigments deposited in developing wing scales (Wittkopp and Beldade, 2009). Melanin production—being involved in a wide range of physiological processes—is conserved across insect orders. For instance, increased melanization in high-altitude moths aids in thermoregulation, resulting in a trade-off between warning coloration and hindwing melanization (Hegna et al., 2013). Melanization is also associated with immunity, wound healing, and protection from both ultraviolet light and parasitoids (Yassine et al., 2012; Bilandžija et al., 2017). Melanin and related pigments—such as DOPA-melanin, dopamine-melanin, NBAD sclerotin, and NADA sclerotin— contribute to the production of black, brown, and yellow coloration (Wright, 1987, Koch et al., 2000; Zhang et al., 2017). The availability of genetic manipulation techniques in non-model systems has allowed elucidation of the role of melanin pathway genes in color adaptations (Zhang and Reed, 2017; Zhang et al., 2017a; Matsuoka and Monteiro, 2018). While the developmental mechanisms underlying pigmentation and patterning are being extensively studied, the evolutionary trajectories of these pigmentation genes remain poorly explored.

We chose six melanin pathway genes—*tan, black, ebony, pale* (*tyrosine hydroxylase*), *arylalkylamine N-acetyltransferase 1* (*aaNAT*) and *DOPA decarboxylase* (*DDC*)—whose developmental function in Lepidoptera coloration is well characterized (Zhang et al., 2017a; Matsuoka and Monteiro, 2018). The functions and phenotypic effects of these genes are summarized in Fig. 1 and Table S1. Melanin synthesis occurs by a branched pathway with tyrosine and uracil as precursors, and different end products (Wright, 1987; Matsuoka and Monteiro, 2018). These precursors are shunted into the melanin pathway by *pale* and *DDC* (Zhang et al., 2017a). *black, ebony*, and *tan* are involved in the synthesis of N-β-alanyl dopamine (NBAD), which is responsible for yellow coloration, while *aaNAT* facilitates the formation of colorless cuticles essential for wing pigmentation (Fig. 1, Table S1). These genes are thus crucial for melanin production and its deposition on Lepidoptera wings. The pigments observed in adults start appearing on the wings and body tissues during late pupal stages, and black melanin is usually the last one to be deposited (Koch et al., 1998; Wittkopp and Beldade, 2009; Matsuoka and Monteiro, 2018). In this study, we trace the molecular evolution of these six genes in 53 species across nine superfamilies of Lepidoptera (total over 100 million years of divergence), and additionally characterize their expression patterns across various developmental stages and tissues of *Papilio polytes* to address the following questions: (a) what selection pressures have shaped the evolution of these genes in the Lepidoptera?, (b) Do these genes show differential activity in sexually dimorphic species?, and (c) Do they play similar roles in larval pigmentation and adult wing patterning and pigmentation?

**Figure 1:**
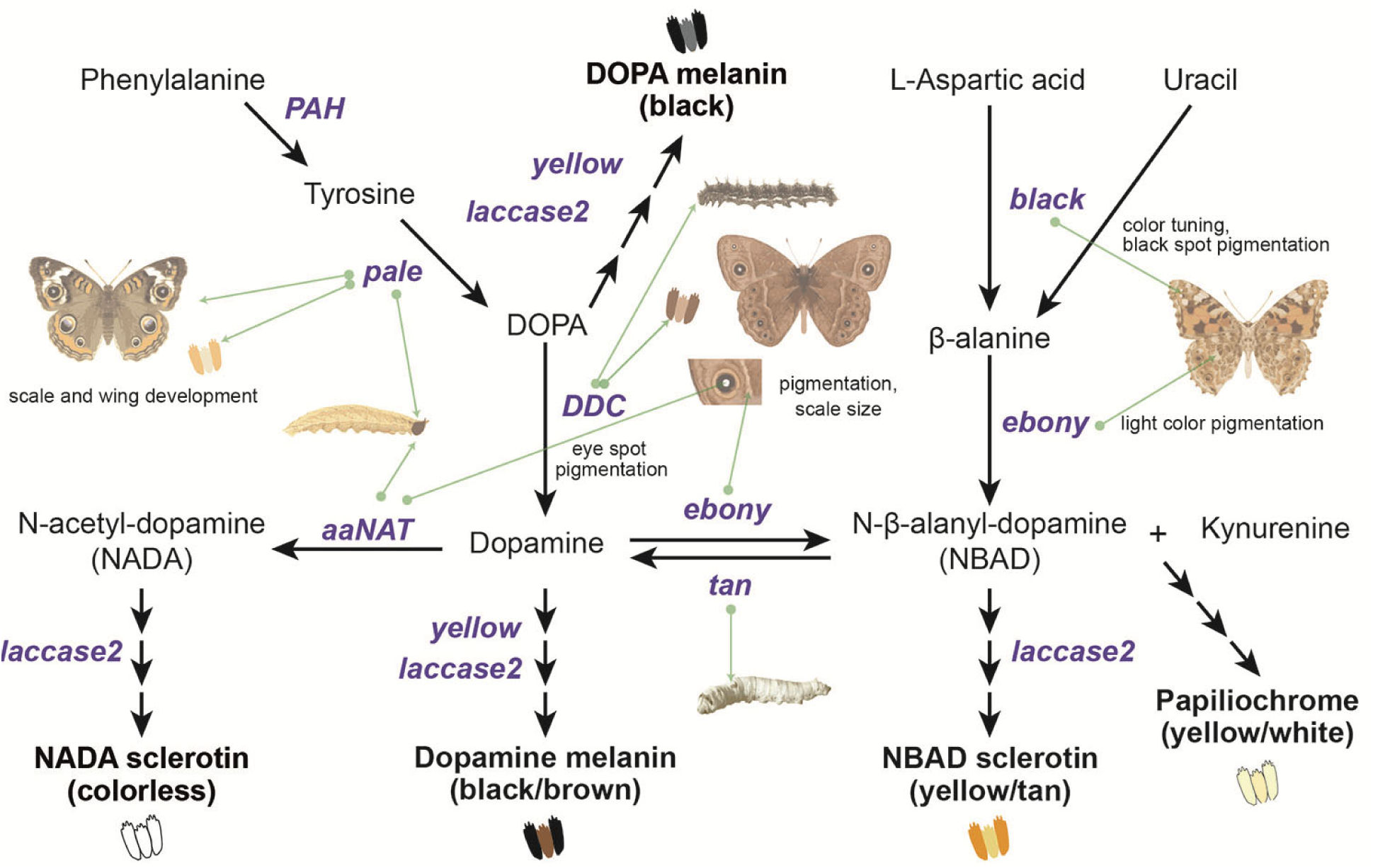
Melanin pathway genes and their phenotypic effects in Lepidoptera. The proposed melanin pathway (Wright, 1987; Zhang et al., 2017) illustrates the genes/enzymes (colored in purple) that catalyze different steps in the production of four major pigments. The end products of the pathway conjugate with sclerotin to produce differently colored scales (Vavricka et al., 2010), as illustrated. Each gene also regulates individual wing elements in different species of butterflies (green arrows), as deciphered from their knockout phenotypes (summarized in Table S1).

## MATERIALS AND METHODS

### Gene sequences and multiple sequence alignment

We downloaded whole genome sequences of 53 species of Lepidoptera from GenBank (http://www.ncbi.nlm.nih.gov), LepBase (http://www.lepbase.org), and GigaDB (http://www.re3data.org), which best represented different families. We selected six genes which are central to pigmentation and patterning pathways namely *tan, black, ebony, pale* (*tyrosine hydroxylase*), *aaNAT* (*Arylalkylamine N-acetyltransferase 1*) and *DDC* (*dopa decarboxylase*) (Wright, 1987; Zhang et al., 2017a; Matsuoka and Monteiro, 2018). We performed exon-wise NCBI local tBLASTn to locate these genes within genomes of the selected species (Table S2). For all the genes, we chose the longest isoform for performing tBLASTn as it incorporates maximum sequence information for the given gene. Using these gene coordinates, we extracted respective genes from the genome using an in-house python script. The downloaded genome sequences showed no evidence of duplication for any of the six melanin pathway genes. Several genome files (such as those of *Bicyclus anynana, Papilio polytes, Papilio xuthus* and *Bombyx mori*) had annotated two copies of *aaNAT* and *black*, but they shared low similarity scores (∼40%) and were therefore not considered as duplicates in our analysis. We performed multiple sequence alignment of each gene with MEGA X muscle aligner with codon alignment option (Kumar et al., 2018). We included homologous gene sequences from Diptera as outgroups for respective genes.

### Gene tree construction and phylogenetic analysis

We constructed a species-level phylogeny with two mitochondrial (*cytochrome c oxidase I, acetyl-CoA acetyltransferase*) and eleven nuclear genes (*elongation factor 1* – *alpha, wingless, ribosomal Protein S5, ribosomal protein S2, isocitrate dehydrogenase, DOPA decarboxylase, glyceraldehyde-3-phosphate dehydrogenase, malate dehydrogenase, catalase, CAD, ribosomal protein S27a / hairy cell leukemia*) that are used commonly in phylogenetic reconstruction (Wahlberg and Wheat, 2008). We performed multiple sequence alignment on these thirteen markers using the codon aligner PRANK v150803 (Löytynoja and Goldman, 2005). For construction of species and gene trees, we calculated the best partition scheme and corresponding sequence evolution model using Partition Finder 2.1.1 (Lanfear et al., 2016). We chose greedy algorithm and Mr. Bayes model of evolution. We selected Bayesian information criterion to compare and choose from the best-fit models. We used a split frequency below 0.01 to assess stationarity and to set the burn-in in Mr. Bayes and then built a consensus tree using the remaining trees (Ronquist et al., 2012).

### Determination of synonymous (dS) and nonsynonymous (dN) substitution rates

To calculate site-wise synonymous and nonsynonymous substitution rates of each gene, we used their respective MUSCLE alignments and the Fixed Effect Likelihood (FEL) method with a maximum-likelihood (ML) approach (Kosakovsky Pond and Frost, 2005). We plotted and compared codon-wise dN, dS, and dN/dS values for each gene (Fig. 2) and performed a Kruskal*-*Wallis test followed by Dunn’s test with Bonferroni correction in R (Derek et al., 2020) We also identified sites that have been subjected to pervasive diversification (dN/dS > 1) or purifying selection (dN/dS < 1) within every gene using FEL. We calculated global omega (R value, which represents dN/dS for the entire sequence) for respective alignments using AnalyzeCodonData function of HyPhy 2.3.14 (Pond et al., 2005; Kosakovsky Pond et al., 2020) (Table 1). To identify conserved domains in the given proteins, we used the Conserved Domain Database (CDD) and the CD-Search Tool (Marchler-Bauer et al., 2012).We separately performed FEL analysis and estimated global omega for these domains to compare gene-wide and domain-wide signatures of selection (Fig. 2 and Table 1).

**Table 1:** Molecular evolution of melanin pathway genes in Lepidoptera. A summary of tests of selection at the levels of individual sites, genes and branches. All the genes studied contained a single conserved domain, except for two domains in *ebony*.

| Gene | *aaNAT* | *black* | *DDC* | *ebony* | *pale* | *tan* |
| --- | --- | --- | --- | --- | --- | --- |
| Number of species | 37 | 44 | 41 | 39 | 38 | 36 |
| Sequence length in nucleotides | 852 | 1545 | 1503 | 2622 | 1689 | 1277 |
| Percentage of sites with pervasive purifying selection | 79.9 | 89.1 | 88.02 | 88.1 | 88.45 | 86.06 |
| Percentage of sites with pervasive diversifying selection | 0 | 0 | 0 | 0 | 0 | 0 |
| Number of branches under episodic diversifying selection | 0 | 2 | 0 | 11 | 2 | 3 |
| Evidence for gene-wide episodic diversifying selection | Yes | Yes | No | Yes | Yes | No |
| Number of sites under episodic selection | 1 | 0 | 0 | 2 | 2 | 3 |
| Global omega value for the gene | 0.09 | 0.06 | 0.04 | 0.08 | 0.04 | 0.08 |
| Omega value for conserved domain 1 | 0.07 | 0.05 | 0.03 | 0.07 | 0.02 | 0.07 |
| Omega value for conserved domain 2 | - | - | - | 0.05 | - | - |

**Figure 2:**
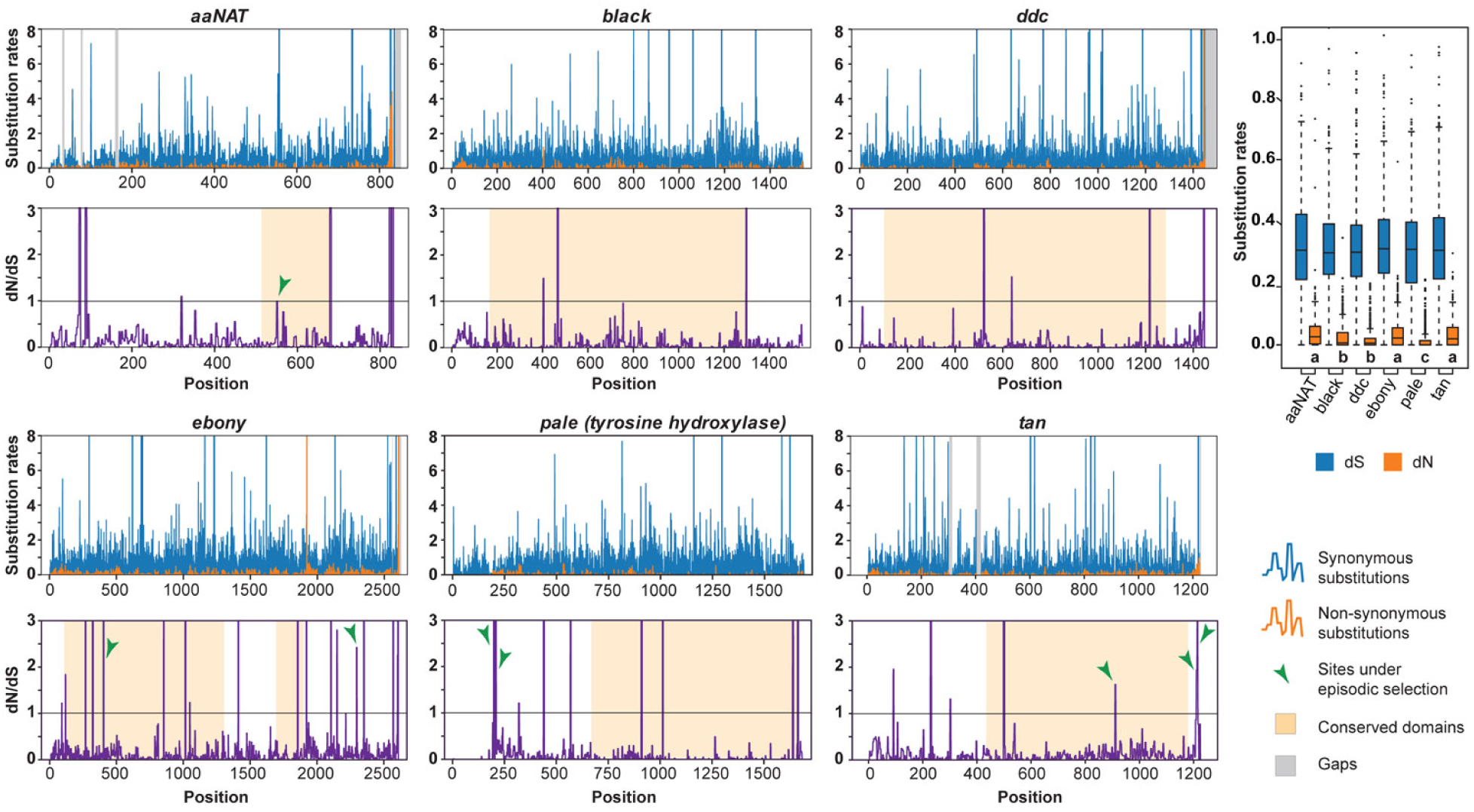
Molecular evolution of selected genes in the melanin pathway. Molecular evolution is shown with individual rates of synonymous (blue) and nonsynonymous substitutions (orange) for each codon (top panels) and dN/dS ratios for individual codons (lower panels). Functional domains for each gene are highlighted along with sites under episodic diversifying selection (green arrows). Rates of substitutions are compared between genes in the boxplot (right) where dN values significantly differ across genes (*p*<0.001, see Table S3). Groups that do not differ statistically are represented by the same letter.

### Branch level, gene-wide and site-wise assessment of molecular evolution

We used aBSREL (adaptive Branch-Site Random Effects Likelihood) method to test for positive selection along each gene tree (Kosakovsky Pond et al., 2011; Smith et al., 2015). We used BUSTED to investigate if any of the six genes have experienced positive selection in at least one site. We carried out MEME (Mixed Effects Model of Evolution) which takes maximum-likelihood approach to test whether individual sites have been subjected to episodic positive or diversifying selection (dN/dS > 1). We performed FEL, MEME, BUSTED and aBSREL on the Datamonkey Adaptive Evolution Server (Weaver et al., 2018), processing the data using in-house scripts.

### Sample collection, RNA extraction, and transcriptome sequencing for quantification of expression

We bred greenhouse populations of *Papilio polytes* from mated wild-caught females. We maintained larvae at 28±4°C on lemon (*Citrus sp*.) and curry plants (*Murraya koenigii*) and adults on Birds Choice™ butterfly nectar. We preserved *Papilio polytes* at different stages of metamorphosis in TRIzol™. We collected eggs at 2, 10 and 24 hours, and at 3 days after oviposition, and pooled five eggs for each sample to get sufficient amount of RNA. We sampled larvae at 1^st^, 3^rd^ and 5^th^ instars. For RNA extraction, we used gutted bodies of 1^st^ and 3^rd^ instar larvae and dissected wing discs from 5^th^ instar larvae and pre-pupae to get three tissue types— forewings, hindwings and gutted body. We collected pupae at 3, 6, and 9 days after pupation, and dissected them to separate the forewings, hindwings, abdomen, thorax, and head. Several pigmentation pathway genes participate in transport and phototransduction in insect eyes (True et al., 2005, Zeigler, 2012). To account for altered sensitivity and visual perception due to the mating status of the butterflies, if any, we collected unmated and mated adults separately and further dissected them to separate abdomen, head, eyes, and thorax. We used pure-breeding mimetic lines for this study. We sampled eggs, 1^st^ and 3^rd^ instar larvae in triplicates, 5^th^ instar larvae, pre-pupae and pupae in duplicates for males and females each, and adults in quadruplets for each sex.

We extracted RNA from preserved tissues using the chloroform-isopropanol-based extraction method. We prepared libraries using TruSeq® RNA Sample Preparation Kit v2 and used Qubit fluorometric quantification. We also used Bioanalyzer to check library profiles. For transcriptome sequencing we used 2×100 PE runs on Illumina HiSeq 2500 and obtained ∼20 million reads for each sample. We performed quality check on these reads using FASTQ, trimmed them using Trimmomatic and aligned them to *P. polytes* reference genome (Nishikawa et al., 2015) using STAR aligner for eggs, larvae and pupae and HISAT2 aligner for adults. We used HTSeq for obtaining raw counts and edgeR pipeline for analyzing and plotting gene expression for all the genes.

## RESULTS

### Melanin pathway genes are single-copy conserved genes under strong purifying selection in Lepidoptera

We reconstructed gene trees for each of the six genes (Fig. S1). Each gene tree was well-supported and it essentially mirrored the species tree in terms of species relationships and divergence, except *aaNAT*, which had poor resolution with a high degree of polytomy. Absence of long branches and broad similarity with the species tree suggest that these genes have evolved with species divergence and do not seem to have undergone rapid evolution that would be incongruent with the species tree in any lineage.

We plotted dN/dS values for each site in each gene (Fig. 2). The molecular evolution of these genes was characterized by predominantly synonymous substitutions, suggesting constrained evolution (dN/dS values for most sites were well below 1, with mean dN/dS value for each gene ranging from 0.04-0.10). Rates for gene-wide synonymous substitutions were similar, but nonsynonymous rates were significantly different between three groups of genes: *black* and *DDC*; *ebony, tan* and *aaNAT;* and *pale* (Dunn’s test, *p*<0.001) (Table S3). Overall, >80% sites showed signatures of pervasive purifying selection (*p*<0.05), but no sites showed signatures of pervasive diversifying selection in any melanin pathway genes (Table 1).

All the genes had a single functional domain except for *ebony*, which contained two (Fig. 2). R values were consistently lower for functional domains compared to the gene-wide values (Table 1). This slight difference between domain-wide and gene-wide R values suggests that these genes are nearly as conserved as their functional domains. In some cases, this may be a result of the domain spanning a large portion of the gene. We observed that the nonsynonymous substitution rate was lowest in *pale*, followed by *black* and *DDC* (Fig. 2, *p*<0.05). Since *DDC, black*, and *pale* are essential for the synthesis of dopamine—a precursor for multiple pathways including neuromodulation, immune functions, and the circadian cycle (Basu and Dasgupta, 2000; Shang et al., 2011; Allen et al., 2011)—the low dN/dS as well as global R values were expected for these genes. On the other hand, *tan, aaNAT* and *ebony* yielded high R values, indicating that they may have evolved under relatively relaxed selection (Fig. 2, Table 1).

### Melanin pathway genes show signs of episodic positive selection

In spite of the overall constrained molecular evolution of the six genes, six sites showed signatures of episodic diversifying selection (*p*<0.05) in *aaNAT, ebony* and *tan*, including some in the otherwise highly conserved domain regions (Fig. 2, Table 1). Interestingly, although *pale* generally shows very constrained molecular evolution similar to *DDC* and *black*, it also had two sites under episodic positive selection. While BUSTED could not detect gene-wide episodic diversifying selection in *DDC* and *tan*, aBSREL found evidence for it in all the genes except *DDC*.

### Melanin pathway genes show stage-specific, often sexually dimorphic expression

We traced the expression of the six genes across different developmental stages, tissues, and sexes in the developmental transcriptome of *Papilio polytes. aaNAT, DDC*, and *pale* showed similar patterns of expression across stages and tissues, as did *black* and *ebony* (Fig. 3). Since *pale, DDC*, and *aaNAT* convert tyrosine to N-acetyl-dopamine in successive reactions with no intermediates (Fig. 1), we expected similar activity patterns of these genes. *aaNAT, DDC* and *pale* showed similar activity across stages with a peak in the pre-pupal stage (Fig. 3), but *pale* additionally showed sexually dimorphic expression at this stage. *DDC* and *pale* expression in most tissues and stages suggests a more general function than melanization-specific activity. Although *ebony* combines the products of *black* and *DDC* to produce NBAD (Fig. 1), its expression pattern was similar to that of *black*, not *DDC*. This suggests that *ebony* activity is more dependent on *black* than on *DDC*. This needs to be experimentally verified. In addition, *ebony* expression also showed sexual dimorphism in pre-pupal stage, while *black* did not.

**Figure 3:**
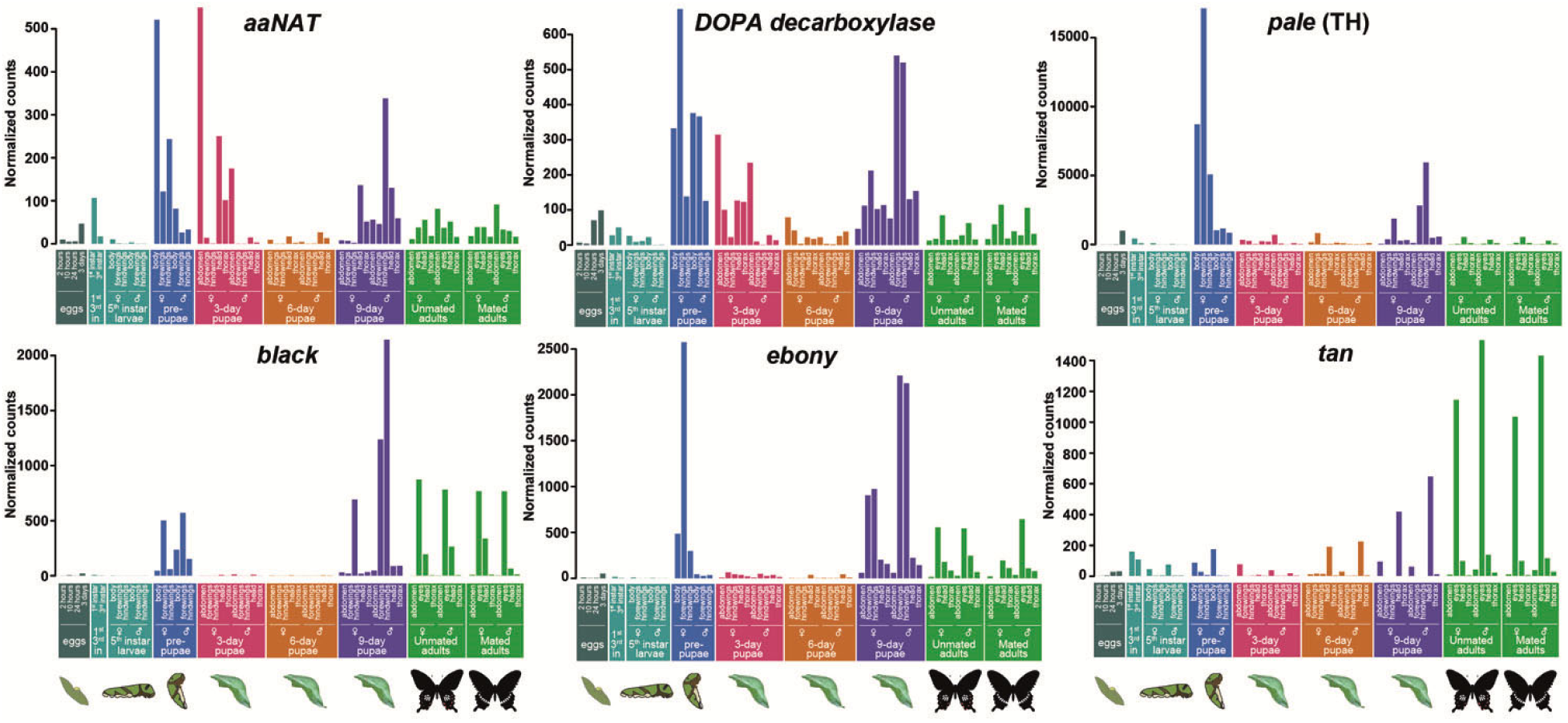
Expression of the six melanin pathway genes across developmental stages of *Papilio polytes*. Normalized counts obtained from transcriptome data are plotted for each stage, tissue and sex (beyond 5^th^ instar larvae). The developmental stages are color-coded and pictorially represented at the bottom.

Unlike any of the other genes, *tan* had a remarkably distinct pattern of expression across development. It showed little to no expression in wings but it was expressed in 1^st^ and 3^rd^ instar larvae, suggesting possible involvement in larval melanization. Expression of *tan, black*, and *ebony* in adult eyes and head may have resulted from their role in photo-transduction. For example, *tan* is involved in metabolism of neurotransmitters in photoreceptors, while *ebony* and *black* play a role in signal transduction in the optic lobe (True et al., 2005, Zeigler, 2012). Because of this, we further tested whether mated and unmated females show differential expression since female visual systems may switch from mate choice to host plant search after mating. However, expression of these genes was similar in unmated and mated females as well as in males.

Finally, four melanin pathway genes (*ebony, pale, aaNAT*, and *DDC*) showed higher expression during larval and early pupal development (pre-pupa and 3-day pupa) in females, but higher expression later in 9-day pupae in males (Fig. 3, Fig. S2). This pattern suggests sex-specific developmental heterochrony.

## DISCUSSION

The melanin pathway is complex and versatile, with products of intermediately placed genes feeding into cross-pathways, performing diverse biological functions (Fig. 1). The feasibility of genetic manipulation in non-model organisms has made it possible to understand the extent of role played by each gene in pigment production in a wide range of organisms (Mazo-Vargas et al., 2017; Zhang et al., 2017a, 2017b; Connahs et al., 2019). We supplement this understanding by studying the molecular evolution of six genes (*tan, black, ebony, pale, aaNAT* and *DDC*) in the melanin pathway that have prominent roles in color production in butterflies and moths (Koch, 1995; Koch et al., 2000a, 2000b; Futahashi and Fujiwara, 2005; Futahashi et al., 2010; Zhang et al., 2017a; Matsuoka and Monteiro, 2018). With each gene governing a generalized or specialized step, the pathway experiences varied constraints at different levels. We estimated synonymous and nonsynonymous substitution rates for each gene using sequence data, and tested for signatures of selection. Our study shows that these six genes are highly conserved, despite 100 million years of evolutionary divergence across the sampled Lepidopteran superfamilies (Misof et al., 2014). However, the degree of conservation varies across genes, and most of them even show some signatures of episodic and positive selection.

Our selection analysis revealed that all the melanin pathway genes studied here show signatures of strong purifying selection. The dearth of nonsynonymous substitutions across these genes indicates a lack of pervasive diversifying selection at a sequence level, although the developmental regulation of these genes produces dissimilar color patterns across species (Wittkopp et al., 2009; Miyazaki et al., 2014; Yassin et al., 2016). While testing for episodic diversifying selection at the levels of individuals sites, branches and genes, *aaNAT, black, ebony, pale* and *tan*, but not *DDC*, showed evidence of diversifying selection. Similarly, we did not find any site-level evidence for diversifying selection in *black*. These differential signatures of selection across the genes are consistent with their estimated nonsynonymous substitutions, which shows that *black* and *DDC* have followed more constrained evolutionary trajectories compared to the other four genes. Our study does not offer insights into the functional roles of these sites under diversifying selection. However, it does point out potential developmental genetic targets for future manipulative experiments. The functional roles of these sites may also become clear using protein structural analysis.

We obtained a few contradictory results in our selection analysis with the three methods used. For example, gene-level analysis using BUSTED inferred the presence of episodic diversifying selection in *black*, but MEME could not detect individual sites under selection. This may be because BUSTED is capable of combining information from weak sites to infer presence of selection at the whole-gene level, unlike MEME, which detects individual sites and might therefore underestimate sites under selection at the gene level. Similarly, we performed branch-level tests of diversifying selection with aBSREL, whose exploratory module is less likely to detect selection compared to targeted testing (Kosakovsky Pond et al., 2011; Smith et al., 2015). This may have resulted in *aaNAT* not showing branch-level selection despite showing site-level and gene-level diversifying selection using other methods.

We explored the spatio-temporal activity of melanin pathway genes in *P. polytes* using a developmental transcriptome dataset. The first four instars of *P. polytes* larvae mimic bird droppings with black, brown, and white coloration on their bodies. We expected to detect activity of melanin pathway genes in the eggs and 1^st^ and 3^rd^ larval instars. However, we did not find substantial larval expression of any of the six genes (except *tan*) even though knockout phenotypes in previous studies have reported a lack of larval pigmentation (Zhang et al., 2017a; Matsuoka and Monteiro, 2018). It is possible that the low expression that we detected suffices in triggering melanization, or we may have missed the transient stage where melanin pathway genes are upregulated in early development. Exploring the expression patterns of other genes in the pathway, such as *laccase2* and *yellow*, may help corroborate either hypothesis. The expression patterns of genes also showed signs of sex-specific developmental heterochrony in melanin production, with expression during pupal development peaking early in females and late in males. While this could be an artefact of sampling, in some cases (especially between male and female pre-pupae and 9-day old pupae), the difference is quite stark. It is possible that males and females differentially invest in melanin production and immunity (or other functions that require melanin pathway intermediates), which would also explain the heterochrony we observed. However, this needs further investigation. With the additional complexity of mimicry in this species, and the use of only mimetic females in this study, the effect of mimicry-related wing-pattern reorganization on melanin pathway genes remains to be explored, and could perhaps be linked to the developmental heterochrony.

Melanin pathway genes such as *tan, ebony* and *yellow* are involved in the evolution of color pattern-related sexual dimorphism and polymorphism (Wittkopp et al., 2009; Miyazaki et al., 2014; Yassin et al., 2016), suggesting that these genes have the potential to evolve novel functions despite evolutionarily constrained sequences. Many butterflies and moths exhibit sexual dimorphism and/or female polymorphism, so they are good systems to study signatures of selection in such clades. This would also require information regarding functional allelic variation and duplicated copies of these genes, which we did not encounter in the species that we studied. Alternatively, examining the regulatory regions of these genes and tracing the molecular evolution of upstream regulators might help detect such signals. We have excluded *yellow* gene family from our study even though substantial work has been done on this in flies and butterflies (Arnoult et al., 2013; Miyazaki et al., 2014; Camino et al., 2015; Zhang et al., 2017a; Matsuoka and Monteiro, 2018). We were unable to identify sequences of this gene family from the downloaded genome sequences with sufficient confidence due to issues with sequence similarity scores. However, the evolution of this gene family has been explored previously (Ferguson et al., 2011).

Our work demonstrates that molecular sequences of the six melanin pathway genes are highly conserved across the vastly diverse Lepidoptera, yet they show some signatures of positive selection. The pattern of conservation and divergence of these genes may be a direct outcome of their functions. We also explored the developmental expression of these genes across tissues and sexes, which recapitulates what we observe in adult stages, but does not provide insights into larval pigmentation. Our work provides a framework in which broad comparisons can be made across genes and species to understand the genetics and evo-devo of complex pathways that produce the remarkable color patterns displayed by insects.

## Supporting information

Table S2: Species list and genome details

## ACKNOWLEDGEMENTS

We thank Atharva Khotpal, Bhavanarayeni R., Sai Guha and Sarvesh Menon for assistance in downloading and analyzing genome sequences; Saurav Baral and Vinod Shankar for help with molecular evolution analysis; Sergei L. Kosakovsky Pond for valuable inputs on HyPhy; NCBS Sequencing Facility for transcriptome sequencing; and NCBS Greenhouse Facility for butterfly breeding facilities.

## AUTHORS CONTRIBUTION

RD and KK designed the study; MK and AP downloaded sequence data and performed molecular evolution analysis; RD performed developmental transcriptome sequencing and analysis; MK and RD prepared figures and wrote the manuscript; KK conceived and directed the project.

## SUPPLEMENTARY MATERIAL

**Table S1:**
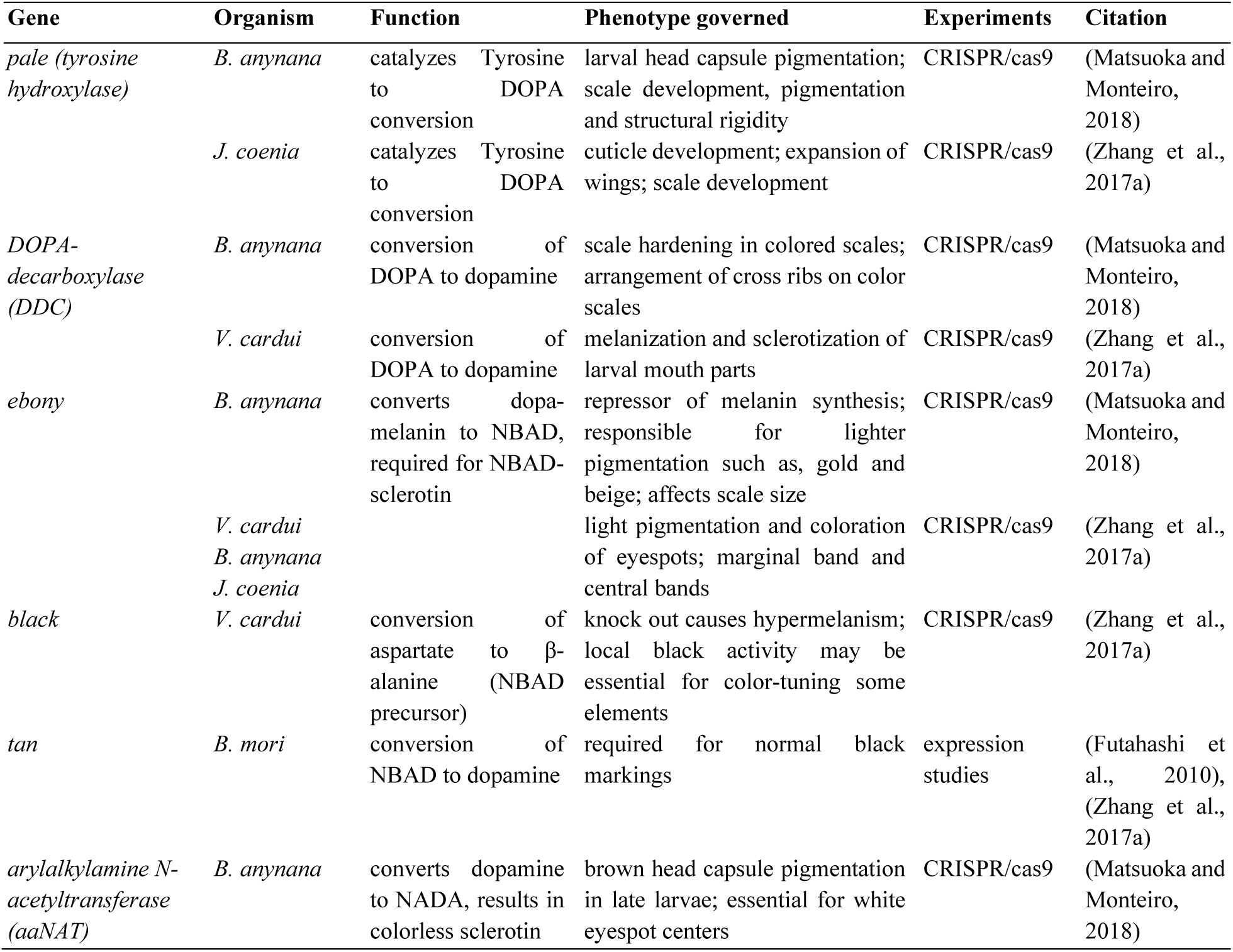
Functional roles of melanin pathway genes selected for this study.

**Table S2: Genomes used in this study.** The accession codes of 53 Lepidopteran genomes and Dipteran outgroups along with details of extracted genes for each comparison are summarized in Excel file TableS2.SpeciesListAndGenomeDetails.xlsx.

**Table S3:** *p*-values for pair-wise comparisons of nonsynonymous substitution rates shown in Fig. 2. Significance of p<0.001 is indicated with ***.

| Gene comparisons for dS | Z statistic | p (unadjusted) | p (adjusted) | Significance |
| --- | --- | --- | --- | --- |
| *aanat-black* | 1.66110505 | 0.09669235 | 1 | - |
| *aanat-ddc* | 1.43518131 | 0.15123544 | 1 | - |
| *black-ddc* | -0.2631673 | 0.79242163 | 1 | - |
| *aanat-ebony* | 0.68234543 | 0.49502055 | 1 | - |
| *black-ebony* | -1.41176586 | 0.15801891 | 1 | - |
| *ddc-ebony* | -1.10020745 | 0.27124175 | 1 | - |
| *aanat-pale* | 1.45306465 | 0.14620581 | 1 | - |
| *black-pale* | -0.28465403 | 0.77590921 | 1 | - |
| *ddc-pale* | -0.01331159 | 0.9893792 | 1 | - |
| *ebony-pale* | 1.12743228 | 0.25955976 | 1 | - |
| *aanat-tan* | 0.66414217 | 0.50659932 | 1 | - |
| *black-tan* | -1.10445915 | 0.26939402 | 1 | - |
| *ddc-tan* | -0.84928716 | 0.39572152 | 1 | - |
| *ebony-tan* | 0.07772577 | 0.9380462 | 1 | - |
| *pale-tan* | -0.85922025 | 0.39021901 | 1 | - |

| Gene comparisons for dN | Z statistic | p (unadjusted) | p (adjusted) | Significance |
| --- | --- | --- | --- | --- |
| *aanat-black* | 5.899968 | 3.64E-09 | 5.45E-08 | *** |
| *aanat-ddc* | 6.949501 | 3.67E-12 | 5.50E-11 | *** |
| *black-ddc* | 1.334072 | 1.82E-01 | 1.00E+00 | - |
| *aanat-ebony* | 1.229317 | 2.19E-01 | 1.00E+00 | - |
| *black-ebony* | -6.543908 | 5.99E-11 | 8.99E-10 | *** |
| *ddc-ebony* | -7.9514 | 1.84E-15 | 2.77E-14 | *** |
| *aanat-pale* | 10.917824 | 9.47E-28 | 1.42E-26 | *** |
| *black-pale* | 6.113977 | 9.72E-10 | 1.46E-08 | *** |
| *ddc-pale* | 4.689634 | 2.74E-06 | 4.11E-05 | *** |
| *ebony-pale* | 13.590144 | 4.58E-42 | 6.87E-41 | *** |
| *aanat-tan* | 2.027325 | 4.26E-02 | 6.39E-01 | - |
| *black-tan* | -4.321267 | 1.55E-05 | 2.33E-04 | *** |
| *ddc-tan* | -5.528088 | 3.24E-08 | 4.86E-07 | *** |
| *ebony-tan* | 1.239223 | 2.15E-01 | 1.00E+00 | - |
| *pale-tan* | -10.077773 | 6.93E-24 | 1.04E-22 | *** |

**Figure S1:**
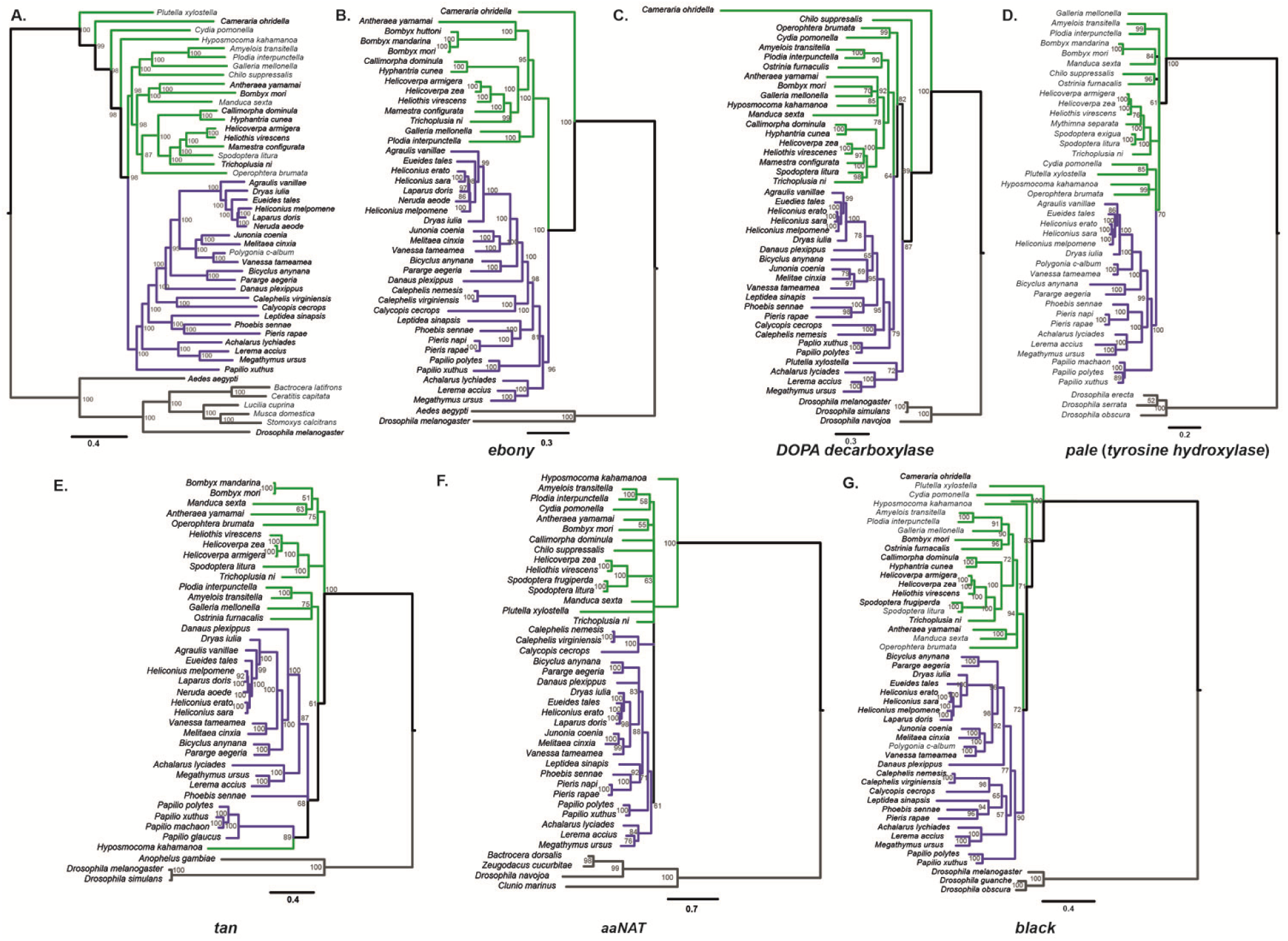
Phylogenetic reconstruction of the species tree and gene trees for the six melanin pathway genes. **A**. Genus level species tree for the taxa used in this study. **B-G**. Gene trees reconstructed using the sequences extracted from genomes of the respective taxa. Green and purple branches represent moths and butterflies, respectively, while gray branches denote outgroups.

**Figure S2:**
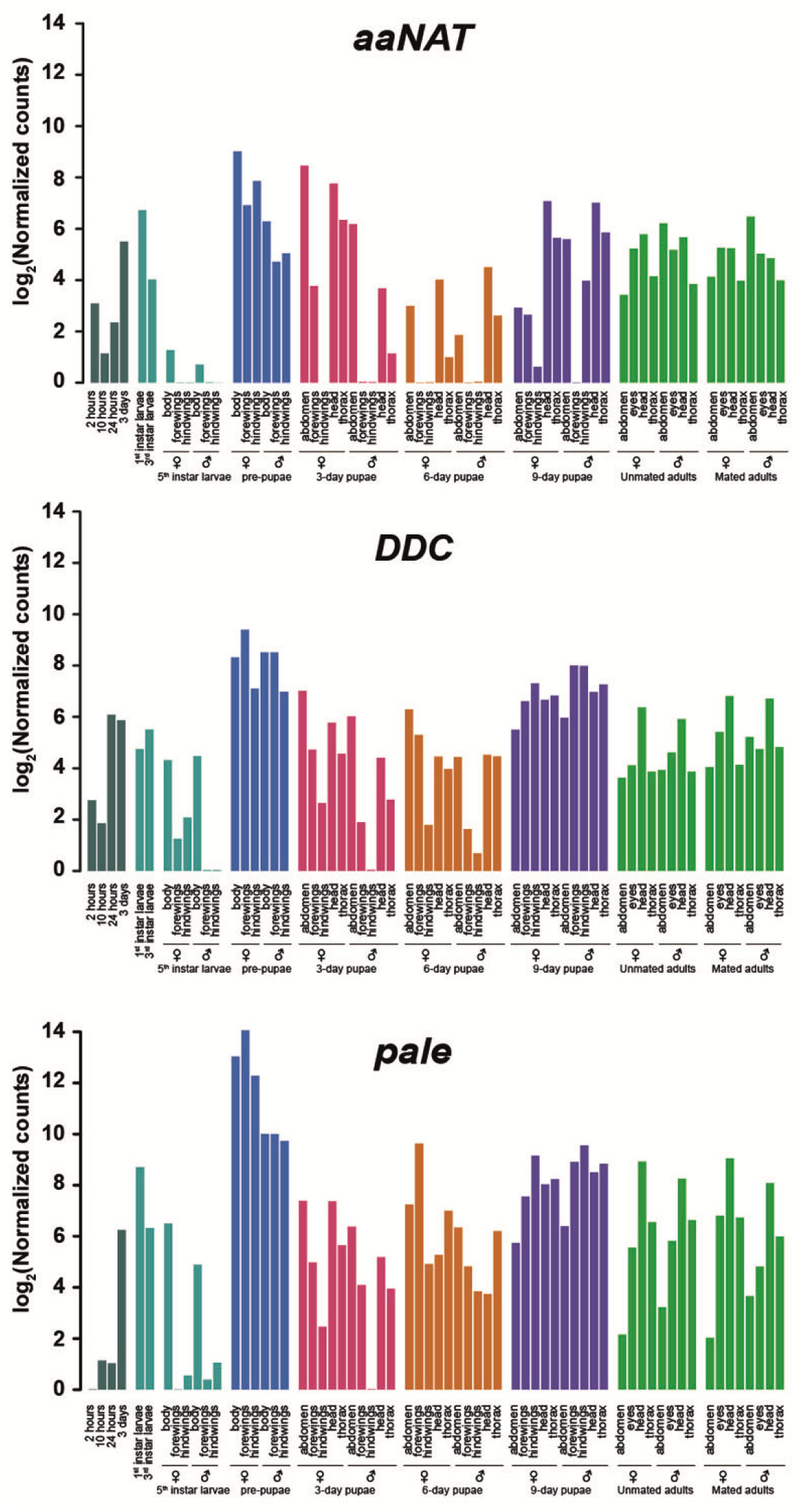
Developmental expression patterns of *aaNAT, DDC and pale* in log2 scale.

